# Characterizing and comparing phylogenetic trait data from their normalized Laplacian spectrum

**DOI:** 10.1101/654087

**Authors:** Eric Lewitus, Leandro Aristide, Helene Morlon

## Abstract

The dissection of the mode and tempo of phenotypic evolution is integral to our understanding of global biodiversity. Our ability to infer patterns of phenotypes across phylogenetic clades is essential to how we infer the macroevolutionary processes governing those patterns. Many methods are already available for fitting models of phenotypic evolution to data. However, there is currently no non-parametric comprehensive framework for characterising and comparing patterns of phenotypic evolution. Here we build on a recently introduced approach for using the phylogenetic spectral density profile to compare and characterize patterns of phylogenetic diversification, in order to provide a framework for non-parametric analysis of phylogenetic trait data. We show how to construct the spectral density profile of trait data on a phylogenetic tree from the normalized graph Laplacian. We demonstrate on simulated data the utility of the spectral density profile to successfully cluster phylogenetic trait data into meaningful groups and to characterise the phenotypic patterning within those groups. We furthermore demonstrate how the spectral density profile is a powerful tool for visualising phenotypic space across traits and for assessing whether distinct trait evolution models are distinguishable on a given empirical phylogeny. We illustrate the approach in two empirical datasets: a comprehensive dataset of traits involved in song, plumage and resource-use in tanagers, and a high-dimensional dataset of endocranial landmarks in New World monkeys. Considering the proliferation of morphometric and molecular data collected across the tree of life, we expect this approach will benefit big data analyses requiring a comprehensive and intuitive framework.

---

Phylogenetic trait data are essential to understanding the evolution of biodiversity. They have been used to identify adaptive radiations (Harmon et al. 2010), infer stabilizing selection (Hansen 1997; Butler and King 2004), measure the phenotypic effects of species interactions (Drury et al. 2018) and environmental fluctuations (Clavel and Morlon 2017), and generally to estimate the role of the phylogeny in how traits evolve over time (Felsenstein, 1973). They are critical to connecting microevolutionary processes of natural selection to macroevolutionary patterns of phenotypic evolution (Hansen and Martins 1996).

A wide range of approaches, reflecting the general interest of trait evolution among evolutionary biologists, have been developed to infer the mode and tempo of phenotypic evolution across clades. These include summary statistics that test for the degree of phylogenetic signal in trait data, such as Blomberg’s *K* (Blomberg et al. 2003), and maximum likelihood-based techniques that fit models to phylogenetic trait data and estimate the rate at which traits evolve (see Pennell and Harmon (2013); Manceau et al. (2016); Lewitus (2018) for a review of currently available models). These models rely on the *a priori* formulation of a phenotypic model, which currently can be reduced to whether traits evolve according to a Brownian process along the phylogeny (Felsenstein 1985), towards a trait optimum (Hansen 1997), as an effect of increasing species diversity (Weir and Mursleen 2013) or environmental fluctuations (Clavel and Morlon 2017), or as a result of interspecific interactions (Drury et al. 2016; Manceau et al. 2016). Insofar as they represent a fixed set of biological scenarios, the reliance on parameterized models ultimately limits our ability to characterize the patterns of trait evolution along a phylogeny and compare those patterns between traits independently of pre-defined evolutionary processes.

In this paper, we introduce an approach for analysing phylogenetic trait data that requires no assumptions about the underlying generative model. This approach allows for comparisons of the evolutionary histories of traits evolving within a phylogenetic clade and the characterization of trait evolution according to an intuitive graph-theoretical system. Our approach is based on the spectrum of the normalized graph Laplacian, which provides a framework for systematically characterizing and comparing the distribution of trait data across a phylogenetic tree. The normalized graph Laplacian has been successfully utilised in the physical sciences to understand how signal processes are embedded within a graph (Shuman et al. 2013) and has been applied to understanding high-dimensional data produced from, for example, social networks (Rohe et al. 2011), text classification (Apté et al. 1994), and image recognition (Zhang and Hancock 2008). It has also begun to be applied to the biological sciences to aid in big data analysis of metabolic networks (Deyasi et al. 2015) and cancer genomics (Rai et al. 2017). More recently, we introduced an approach for comparing and characterising phylogenies (Lewitus and Morlon 2016a) using the spectral density profile of the graph Laplacian of the distance matrix of a phylogeny, the so-called modified graph Laplacian (MGL), which is able to infer diversification patterns within a phylogeny, as well as directly compare patterns between phylogenies, absent any *a priori* model specification (Lewitus and Morlon 2016b). Together, these applications show the strength of applying the graph Laplacian. However, despite its widespread utility, no such framework has been developed for characterizing and comparing phylogenetic trait data.

We first describe how to construct the spectral density profile of the normalized graph Laplacian for phylogenetic trait data and demonstrate how to interpret it in terms of specific properties of phenotypic evolution. We use simulations to show how the profiles relate to conventional metrics of phylogenetic signal and models of trait evolution. We show how to compute the distance between profiles and cluster phylogenetic trait data based on those distances. Finally, we illustrate the utility of this approach for assessing whether distinct trait evolution models are distinguishable using the Cetacean phylogeny. We also illustrate the application of the approach on functional trait data for tanagers (*Thraupidae*) and geometric morphometric data for the endocrania of New World monkeys (*Platyrrhini*). We think that such a non-parametric and comprehensive framework for studying phylogenetic trait diversification will be a valuable complement to existing model-based approaches.

## Materials and Methods

### Implementation

Below, we describe how to use the normalized modified graph Laplacian (nMGL) to construct a spectral density profile for traits (i.e., unidimensional continuous extant tip data) on a phylogeny, how to characterize the profile in terms of evolutionary patterning, and how to compute the distance between profiles. We implemented these functionalities in the R package *RPANDA* freely available on CRAN (Morlon et al. 2016). In the analyses detailed below, phylogenies were simulated using the *R* package *TESS* (Höhna 2013); trait data for BM, OU and ACDC models were simulated using *mvMORPH* (mvSIM function Clavel et al. (2015)) and for DD and MC models with *RPANDA* (sim_t_comp function). Blombergs K was computed using *phytools* (Revell 2012); and MDI was computed using *geiger* (Harmon et al. 2008).

### Construction of the Spectral Density Profile for Phylogenetic Trait Data

We aim to provide a non-parameteric framework for characterizing and comparing patterns of phylogenetic traits (i.e., tip data) for a given phylogeny. We consider a fully bifurcated tree composed of *m* terminal branches (Fig. 1A). We note 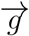 a vector of unidimensional continuous extant trait data associated to this tree. We consider this data as a particular kind of graph, *G* = (*N,E,w*), composed of nodes representing extant species, edges delineating the relationships between nodes, and a weight associated to each edge, computed as *w*(*i,j*) = *d_i,j_*|*g_i_* – *g_j_*| where *d_i,j_* is the phylogenetic distance between tips *i* and *j* and *g_i_* is the trait value at tip *i*. Hence, the weight is a combination of phylogenetic and trait distances between two extant species. In Lewitus and Morlon (2016a), the nodes in the graph represent both extant species and internal splitting events in the phylogeny; here we limit the nodes to extant species, as internal splitting events do not have associated trait data. We consider Θ the matrix of weights (Fig. 1B) and D the degree matrix (the diagonal matrix where diagonal element i is computed as 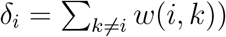. We construct the normalized modified graph Laplacian (nMGL, see Table 1), defined as *D*^−1/2^(*D* – Θ)*D*^−1/2^, which is distinguished from the non-normalized graph Laplacian (*D* – 0) because it is normalized by D. While the normalized version of the graph Laplacian loses some information on the size of the graph compared to the non-normalized version, it is more sensitive to fine-scale features of the graph (Banerjee and Jost 2008). Our approach aims to characterize and compare traits on the same phylogenetic tree (rather than traits between different phylogenetic trees) and so the size of the graph (i.e., of the tree) is not important. The nMGL is a *m* x *m* positive semi-definite matrix. It therefore has n non-negative eigenvalues, _*n*_λ_1_ ≥ _*n*_λ_2_ ≥ ≥ _*n*_λ_*m*_ ≥ 0 (throughout, the n subscript preceding symbols highlights that we are considering the normalized graph Laplacian). We convolve them with a Gaussian kernel to ensure a continuous distribution (Banerjee and Jost 2008). The spectral density profile (SDP) of _*n*_λ from the nMGL, defined as 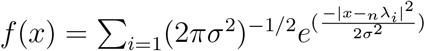, is plotted as a function of _*n*_λ as 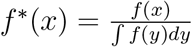 (Fig. 1C). Considering the success of previous work showing the capacity of spectral density profiling for differentiating graphs generated by different processes (Banerjee and Jost 2009; Arenas et al. 2006; McGraw and Menzinger 2008; Lewitus and Morlon 2016b), and particularly the framework we recently introduced for characterizing and comparing phylogenies based on their spectral density profiles (Lewitus and Morlon 2016a), we hypothesized that the spectral density profile of the nMGL would be a powerful tool for characterizing and comparing trait evolution within phylogenetic clades.

**Figure 1:**
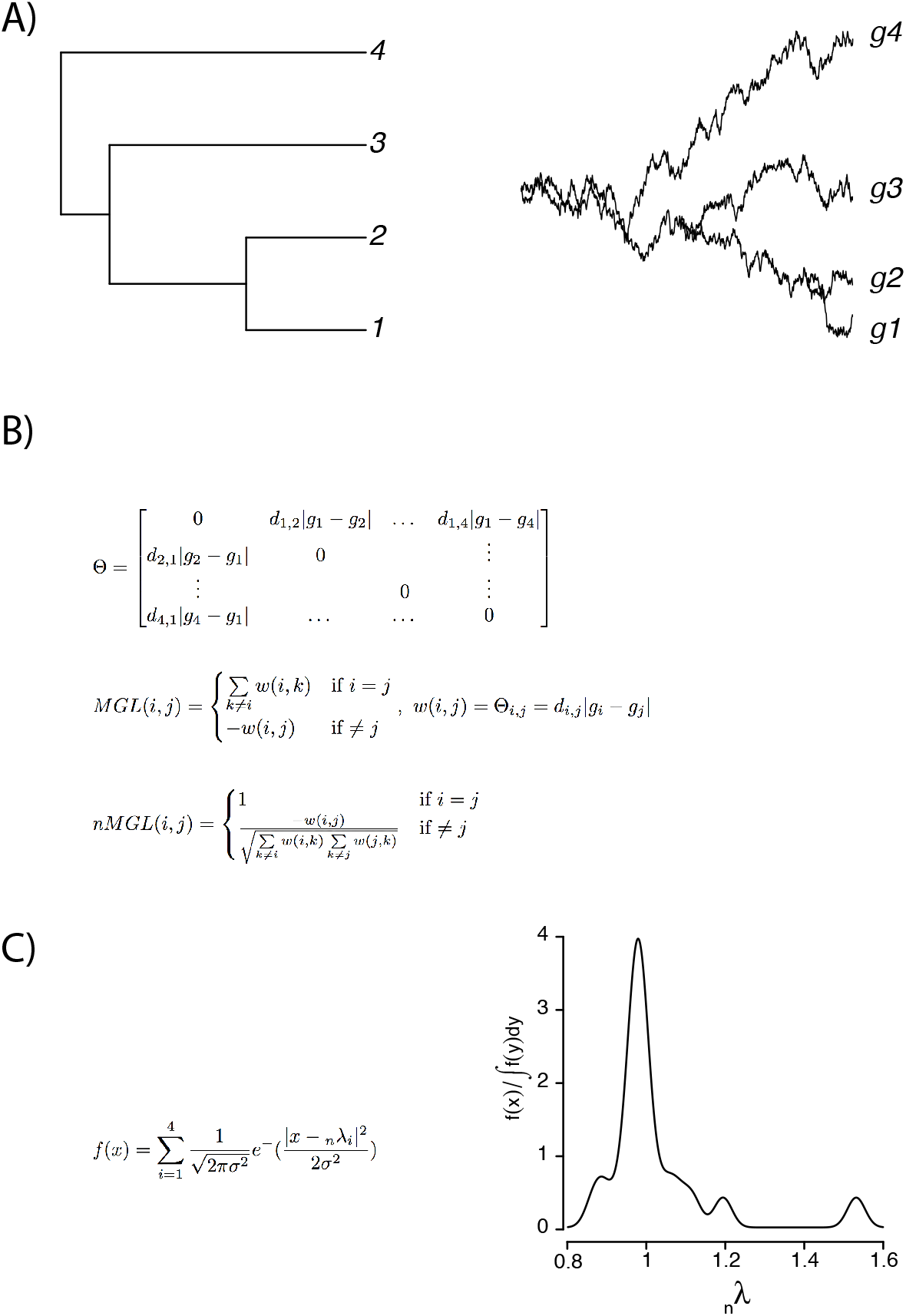
Pipeline for constructing the spectral density profile for the nMGL of phylogenetic trait data. (A) Given a phylogenetic tree with *m* terminal branches and unidimensional, continuous, extant trait data for *m* tips, (B) take the Hadamard product of the difference matrix of the trait data (|*g_i_* – *g*|) and the matrix of phylogenetic branch-lengths between tips (*d_i,j_*), such that Θ = *d_i,j_*|*g_i_* – *g_j_*| at *i* ≠ *j* and zero along the diagonal. The weighted MGL, *D* – Θ, where *D* is the degree matrix of Θ, is computed as the weighted value of (*i, j*), –Θ(*i, j*) = −*w*(*i, j*), at *i* ≠ *j* and as 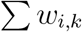 for *i* = *j*. The normalized MGL (nMGL) is normalized by *D*, so that nMGL= *D*^−1/2^(*D* – Θ)*D*^−1/2^, resulting in unity along the diagonal and negative the weighted value of (*i, j*) divided by the square-root of the product of *δ_i_* and *δ_j_* for *i* = *j*. (C) The spectral density is obtained by convolving the eigenvalues, _*n*_λ, computed from the nMGL with a Gaussian kernel and then plotting the density of _*n*_λ.

**Table 1:** Glossary of graphical and statistical terms.

| *symbol* | *descriptor* | *significance* | *value* |
| --- | --- | --- | --- |
| nMGL | normalized modified graph Laplacian | the distance matrix of the phylogeny weighted by the differences in trait values between tips and normalized by the degree matrix | positive semi-definite symmetric matrix |
| $\lambda_n$ | eigenvalues | eigenvalue calculated from the nMGL | $\sim 1 < \lambda_n \leq 2$ |
| SDP | spectral density profile of the nMGL | the density profile of eigenvalues calculated from the nMGL | kernel density estimate of $\lambda_n$ |
| $\lambda_n^*$ | splitter | the maximum eigenvalue; reflects bipartiteness (i.e., monophyletic clustering of trait data) | $1 < \lambda_n^* \leq 2$ |
| $\psi_n$ | fragmenter | the skewness of the SDP; reflects discreteness | $0 < \psi_n < \infty$ |
| $\eta_n$ | tracer | the maximum height of the SDP; reflects phylogenetic signal | $0 < \eta_n < \infty$ |

### Interpreting Spectral Density Profiles for the nMGL

The spectrum of _*n*_λ computed from the nMGL represents primarily global properties of the structure of trait evolution within a phylogenetic clade. Each _*n*_λ reflects the connectivity (in terms of edge-length) and difference in trait value between one tip and all other tips in a phylogeny. We know from the substantial body of existing work on the normalized graph Laplacian that large _*n*_λ are characteristic of sparse neighbourhoods typical of highly divergent terminal branches (both in terms of trait value and phylogenetic distance) and small _*n*_λ are characteristic of denser neighbourhoods typical of barely divergent terminal branches (Chung 1996; Chen et al. 2004). Additionally, for the normalized graph Laplacian, 0 ≤ _*n*_λ ≤ 2 (Bauer and Jost 2009), which in the case of dense matrices (i.e., no zero entries, like 0) becomes ~ 1 ≤_*n*_λ≤ 2 (Banerjee and Jost 2009). Therefore, as trait differences between closely related tips become smaller, _*n*_λ ≃ 1 accumulate and, as trait differences between closely related tips become larger, _*n*_λ > 1 accumulate. Importantly, trait differences here are relative, so small (or large) trait differences are only small (or large) in regard to the distribution of trait differences across the tree. Also, because the weights used to compute the nMGL are products of phylogenetic and trait distances, it is impossible to separate the relative contribution of each of these distances on the SDP. This is one of the reasons why the nMGL is useful for comparing various trait distributions on a given fixed tree (with fixed phylogenetic distance) and not across different trees.

We define three summary statistics computed from the spectrum of _*n*_λ – the tracer, the fragmenter, and the splitter – that together define the phylogenetic trait space. Traits evolved under different evolutionary scenarios on the same tree occupy different regions of this space (Fig. 2).

**Figure 2:**
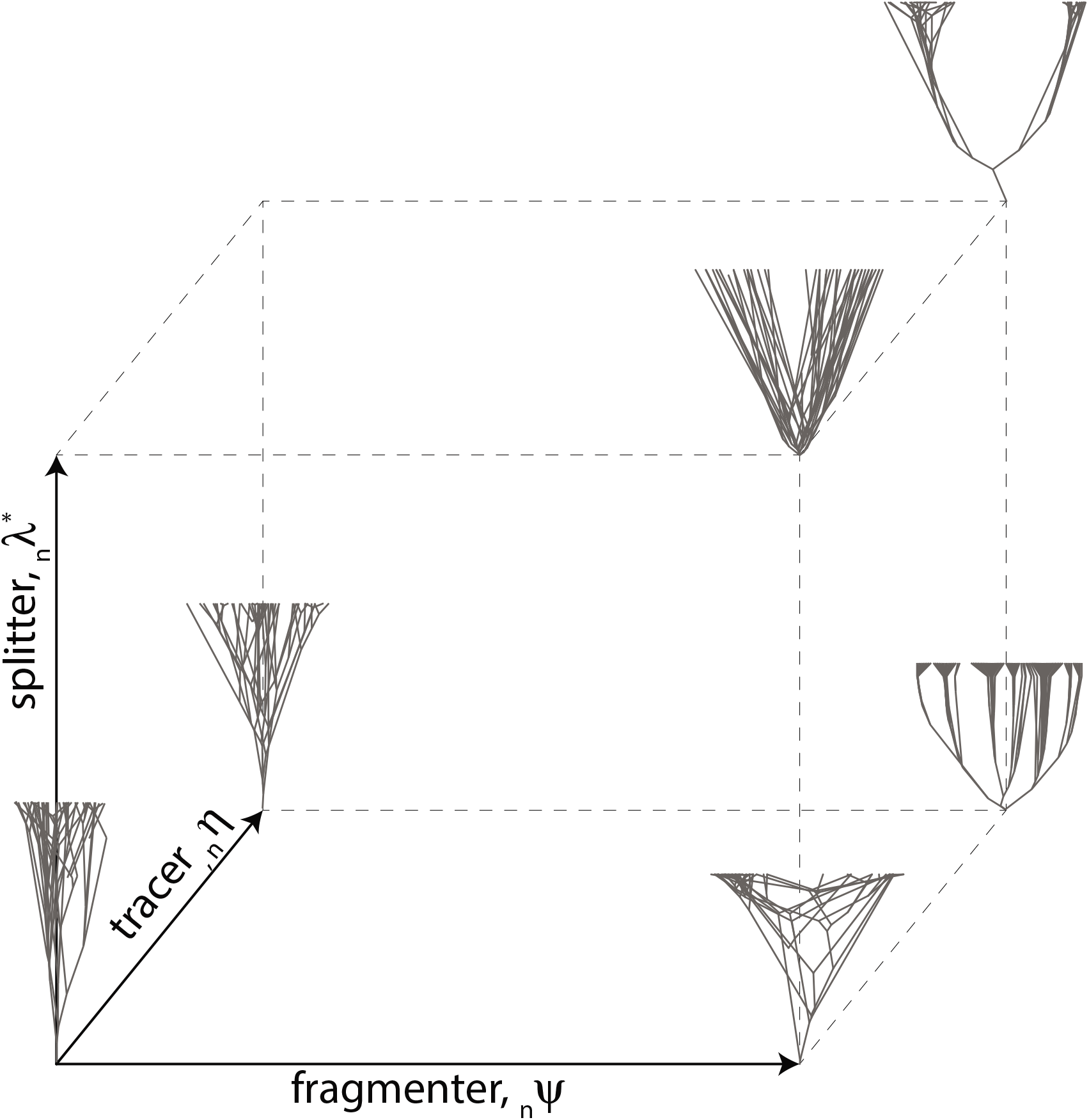
Defining phylogenetic trait space. Any phylogenetic trait data can be placed in a three-dimensional space defined by the splitter (_*n*_λ^*^), the fragmenter (*_n_ψ*), and the tracer (*_n_η*), which broadly represent the bipartiteness, discreteness, and phylogenetic signal, respectively, of the phylogenetic trait data. Hypothetical traitgrams are placed in the corners of the space, illustrating the type of patterns expected in those corners. Traitgrams are generated on the same phylogenetic tree under different trait evolution parameters.

**The tracer** is the peak height of the SDP, denoted *_n_η*, and computed as the ln-transformed maximum value of *f*^*^(*x*); it represents the iteration of _*n*_λ around a single value. Higher tracer values mean smaller differences between closely related tips (low within-clade variance) and larger differences between distantly related tips (high among-clade variance). Therefore, we expect the tracer to be a good measure of phylogenetic signal. In order to test this, we compared the tracer to conventional estimates of phylogenetic signal on trait data simulated on a phylogenetic tree. We simulated a single birth-death tree with 100 tips at 20 million years under constant speciation (0.2) and extinction (0.05) rates (throughout, rates of speciation and extinction are expressed in event per lineage per million year) and simulated 500 trait datasets on that tree under a Brownian motion (BM) model of trait evolution with variance *σ*^2^ = 0.01 (Cavalli-Sforza and Edwards 1967), an exponentially accelerating (AC) model with rate value *β* = 1.5 and *σ*^2^ = 0.01, an exponentially decelerating (DC) model with rate value *β* = −0.1 and *σ*^2^ = 0.01 (Blomberg et al. 2003; Harmon et al. 2010), and a white-noise model by randomly drawing trait values from a normal distribution (with a mean of zero and standard deviation of one). For each of the three first models, we set the root value at 0. For each dataset, we estimated Blomberg’s *K*, which measures the partitioning of variance using a BM model as reference, where *K* > 1 means close relatives resemble one another more than expected under BM, and *K* < 1 means they resemble one another less (Blomberg et al. 2003), and the morphology disparity index (MDI), which is a measure of the difference between the observed diversity through time curve and that expected under a BM model, where a higher MDI indicates that higher subclade disparity than expected under a BM model (Foote 1997; Harmon et al. 2003; Slater et al. 2010). We fit OLS regression models between *_n_η* and both Blomberg’s *K* and MDI for the 500 trait datasets.

**The fragmenter** is the skewness of the SDP, denoted *_n_ψ*, and computed as the ln-transformed 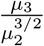, where *μ_i_* is the ordinary *ith* moment of the distribution; it represents the relative abundance of small and large _*n*_λ. Therefore, as trait space becomes more clustered, irrespective of phylogenetic signal, the proportion of small _*n*_λ increases and so does the fragmenter. Therefore, we expect the fragmenter to be a good measure of the discreteness of trait space. In order to test whether the fragmenter captures discrete clusters of extant trait data, we simulated a single birth-death tree with 200 tips at 20 million years under constant speciation (0.2) and extinction (0.05) rates and simulated 200 datasets of discrete trait space under low and high phylogenetic signal. For low phylogenetic signal, we simulated trait data on four macroevolutionary landscapes (Boucher et al. 2017), each defined by a different polynomial function: *V*(*x*) = *x*^2^, *V*(*x*) = *x*^4^ – 0.5*x*^2^, *V*(*x*) = *x*^6^ – 0.5*x*^2^, and *V*(*x*) = *x*^8^ – 0.5*x*^2^, where the landscape is estimated as *e*^−*V*(*x*)^. Here, an increase in the exponent of the first term generates a more discretized trait distribution (i.e., a deeper well in the macroevolutionary landscape). We set *σ*^2^ = 0.5, the root value equal to 5, and the trait boundaries at [0, 10]. We plotted the landscapes as defined by the polynomial functions, histograms of the trait data for each landscape as realized in the simulations, and the spectral density profiles of each dataset. For high phylogenetic signal, we simulated trait data on the same birth-death tree under a DC model with rate value *β* = −0.6, −0.3, 0 and *σ*^2^ = 0.1, where more negative values of *β* indicate more decelerated rates (Blomberg et al. 2003; Harmon et al. 2010). This generated trait data distributed in discrete monophyletic clusters across the tree. For the low and high phylogenetic signal datasets, we computed the fragmenter and compared values as a function of macroevolutionary landscape and of *β*.

**The splitter** is the principal _*n*_λ, denoted (_*n*_λ^*^); it is diagnostic of the disjointedness of a graph, where larger splitter values imply a more bipartite structure (Banerjee and Jost 2008; Bauer and Jost 2009). In macroevolutionary terms, as traits become increasingly bimodally distributed with a strong phylogenetic signal, the splitter increases. As _*n*_λ ≤ 2 for the nMGL, the splitter≃2 when a clade is composed of two phylogenetically distinct subclades with different mean trait values. To assess the relationship between the spectral density profile and differences in mean trait values on a phylogeny, we simulated a single birth-death tree with constant speciation (0.2) and extinction (0.05) rates with 200 tips at 20 million years. We then simulated BM models (*σ* = 0.01) with *q* differences in mean trait values for *q* = 0 – 4 by defining different mean trait values for *q*+1 monophyletic sets of tips, where the mean trait value for *q*_0_ was randomly drawn from a normal distribution with a mean value between 0 – 1 (and standard deviation of one) and subsequent mean trait values were defined as two-times the previous mean. We then compared _*n*_λ^*^ for each set. The value of the splitter is expected to correlate with the disjointedness of the graph, where higher values indicate the nMGL is more bipartite and so can be segregated into two monophyletic groups with distinct mean trait values (Bauer and Jost 2009). To test whether there were, in fact, two monophyletic clusters, we used k-means clustering (for k=2) on the nMGL of the phylogenetic trait data. We then calculated the average branch-length distance between tips in cluster 1 and tips in cluster 2. For phylogenetic trait data that can be separated into two monophyletic clusters, the average between-group distance will equal two times the crown age of the tree. We present this as a heuristic test of the monophyly of trait values when the splitter ≃ 2. To test the effect of phylogenetic signal on the splitter value, we simulated 10 trait datasets with one difference in mean trait values on a 100-tip constant-rate birth-death tree as above. We then randomized the distribution of tip data within each cluster 100 times and compared the resulting splitter value for the randomized trees against the original splitter value. We compared the splitter values for the two-cluster BM datasets and the randomized two-cluster datasets to 100 datasets simulated under a simple BM process (with no clusters and *σ* = 0.01).

To test the effect of erroneous data on the nMGL, we simulated trait data under a BM process on a 100-tip constant-rate birth-death tree (*σ* = 0.01). We tested both the effect of increasing the amount of error and increasing the number of tips with error. For the former, we introduced error on 10% of randomly drawn tips as a sampling variance equal to n times the standard error for *n* = 1, 2, 3. For the latter, we introduced error on 20, 30, 40,100% of tips as a sampling variance equal to the standard error. We simulated 100 datasets for each scenario. We compared the resulting splitter, tracer, and fragmenter values to BM datasets (*σ* = 0.01) and ACDC datasets (*β* = −1.1, *σ* = 0.01) simulated on the same tree and with no introduced error.

### Clustering nMGLs from Their Spectral Density Profiles

To demonstrate whether we can distinguish phylogenetic trait data simulated under trait models we know are distinguishable, we clustered nMGLs constructed for trait data on the same phylogeny under different trait models. To cluster nMGLs, we computed the Jensen-Shannon distance between spectral density profiles. The Jensen-Shannon distance is defined as

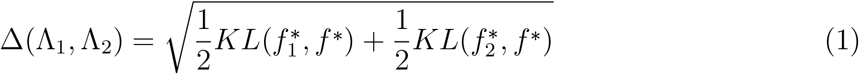

where 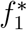 and 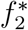 are spectral densities for profiles 1 and 2, 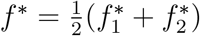, and KL is the Kullback-Leibler divergence measure for the probability distribution (Endres and Schindelin 2003). We then cluster the matrix of Jensen-Shannon distances for each profile pair using hierarchical clustering with boostrap resampling and k-medoids clustering using optimal silhouette width, *s*(*i*), which is a measure of the between/within-variance of each datapoint *i* assigned to a cluster; data are typically considered structured at 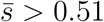 (Szekely and Rizzo 2005; Reynolds et al. 2006). In each case, the number of clusters is not set *a priori*.

We tested the efficacy of clustering on profiles using trait datasets simulated on birth-death trees. We simulated a total of 1500 trait datasets under a BM model of trait evolution with variance *σ*^2^ = 0.01 (Cavalli-Sforza and Edwards 1967), an exponentially AC model with rate value *β* = 1.5 and *σ*^2^ = 0.01, and an exponentially DC model with rate value *β* = –0.1 and *σ*^2^ = 0.01 (Blomberg et al. 2003; Harmon et al. 2010). For each model, we set the root value at 0. We visualised this clustering by plotting the profiles in a multidimensional space defined by _*n*_λ^*^, *_n_ψ*, and *_n_η*.

We tested the ability of the spectral density profile to find meaningful clusters of trait models on different tree shapes and sizes. To test for the effect of tree shape, we simulated trait datasets on 200-tip birth-death trees (with a max age of 20Ma) with constant speciation (0.2) and extinction (0.02) rates, with decreasing speciation (0.1 * *e*^−0.2*t*^) and constant extinction (0.02) rates, and with increasing speciation (0.1 * *e*^0.1*t*^) and constant extinction (0.05) rates. We conducted analyses on identical trees without pruning extinct lineages, resulting therefore in non-ultrametric trees, to test whether the profiles of different models were more distinguishable on a non-ultrametric tree compared to an ultrametric tree, as is expected from likelihood-based approaches (Cooper et al. 2015). To test for the effect of tree size, we simulated 6000 trait datasets under the same BM, AC, and DC trait model parameters on birth-death trees with constant speciation (0.2) and extinction (0.05) rates with 20, 50, 100, 200, and 500 tips (with a max age of 20Ma). As above, phylogenies were simulated using the *R* package *TESS* (Höhna 2013) and trait data were simulated using mvMORPH (Clavel et al. 2015).

### Applications

To illustrate our approach, we demonstrate three applications. First, we used the Cetacean phylogeny (87 spp.) (Steeman et al. 2009) to illustrate how our approach can be used to assess the distinguishability of different trait evolution models in a particular clade. We simulated six trait models under a range of parameter values on the Cetacean phylogeny: BM with *σ*^2^ = 0.1 – 5; Ornstein-Uhlenbeck (OU) with strength of pull towards an optimum *α* = 1 – 20 and *σ*^2^ = 0.1; exponential diversity-dependence (DD) with slope parameter *r* = −1.1 – −0.1 and *σ*^2^ = 0.1; AC with rate value *β* = 1.1 – 1.5 and *σ*^2^ = 0.1; DC with rate value *β* = −0.5 – −0.1 and *σ*^2^ = 0.1; and matching competition (MC) with the strength of competition *S* = −1.1 – −0.1 and *ς*^2^ = 0.1. For each model, we simulated 500 datasets with the root value set to zero. For all datasets, we computed the spectral density profile and clustered them using hierarchical and k-medoid clustering. Second, we used a tanager phylogeny (350 spp.) (*Thraupidae*) with 27 phylogenetically corrected principal component traits (pPC traits) spanning traits related to song, plumage, and resource-use taken from Drury et al. (2018). Ideally, we would have used non-phylogenetically corrected PCs but these were not available. We computed the spectral density profiles for the pPC traits and clustered them using hierarchical and k-medoid clustering and computed their spectral density profile summary statistics. Finally, we used a geometric morphometrics dataset consisting of 399 three-dimensional Procrustes superimposed landmark coordinates describing the external brain shape of 48 species of New World monkeys (*Platyrrhini*) (Aristide et al. 2016). For each landmark, we computed the Euclidean distance between the landmark and the clade mean for that landmark, in order to reduce the dimensionality of the data. We refer to these distances simply as landmarks. We computed the spectral density profile for each of the 399 landmarks and clustered them using hierarchical and k-medoid clustering and plotted their spectral density profile summary statistics in multidimensional space. In order to test how much information was lost in this dimensionality reduction (Monteiro et al. 2000; Uyeda et al. 2015), we also clustered the profiles computed separately for the coordinates along each axis. Even though these axes may not necessarily be aligned with the most biologically meaningful directions of variation, it is a straightforward and convenient way of analyzing the data.

## Results

### Interpreting the Spectral Density Profile for Phylogenetic Trait Data

The shape of the spectral density profile of the nMGL reveals many aspects characteristic of the underlying evolution of a trait within a phylogenetic clade. Specifically, the tracer (peak height, *_n_η*), the fragmenter (skewness, *_n_ψ*) and the splitter (principal _*n*_λ, _*n*_λ^*^), of the profile may be interpreted in terms of the evolutionary history of the trait (Fig. 2).

The tracer summary statistic represents the peak height of the spectral density profile. In macroevolutionary terms, this is indicative of the phylogenetic signal of a trait, where larger *_n_η* indicate more phylogenetic signal (Fig. 3A-C). We show that the tracer is strongly correlated with conventional summary statistics of phylogenetic signal, with *_n_η* increasing with Blomberg’s *K* (*y* = 3.44 – 4.13*x* + 1.36*x*^2^, *R*^2^ = 0.96, *P* < 0.01) and decreasing with MDI (*y* = −2.65 + 2.23*x* – 0.34*x*^2^, *R*^2^ = 0.93, *P* < 0.01) (Fig. 3D). White-noise models fall at the lowest end of tracer values, converging with AC models simulated with *β* = 1.5 in terms of tracer values (Fig. 3).

**Figure 3:**
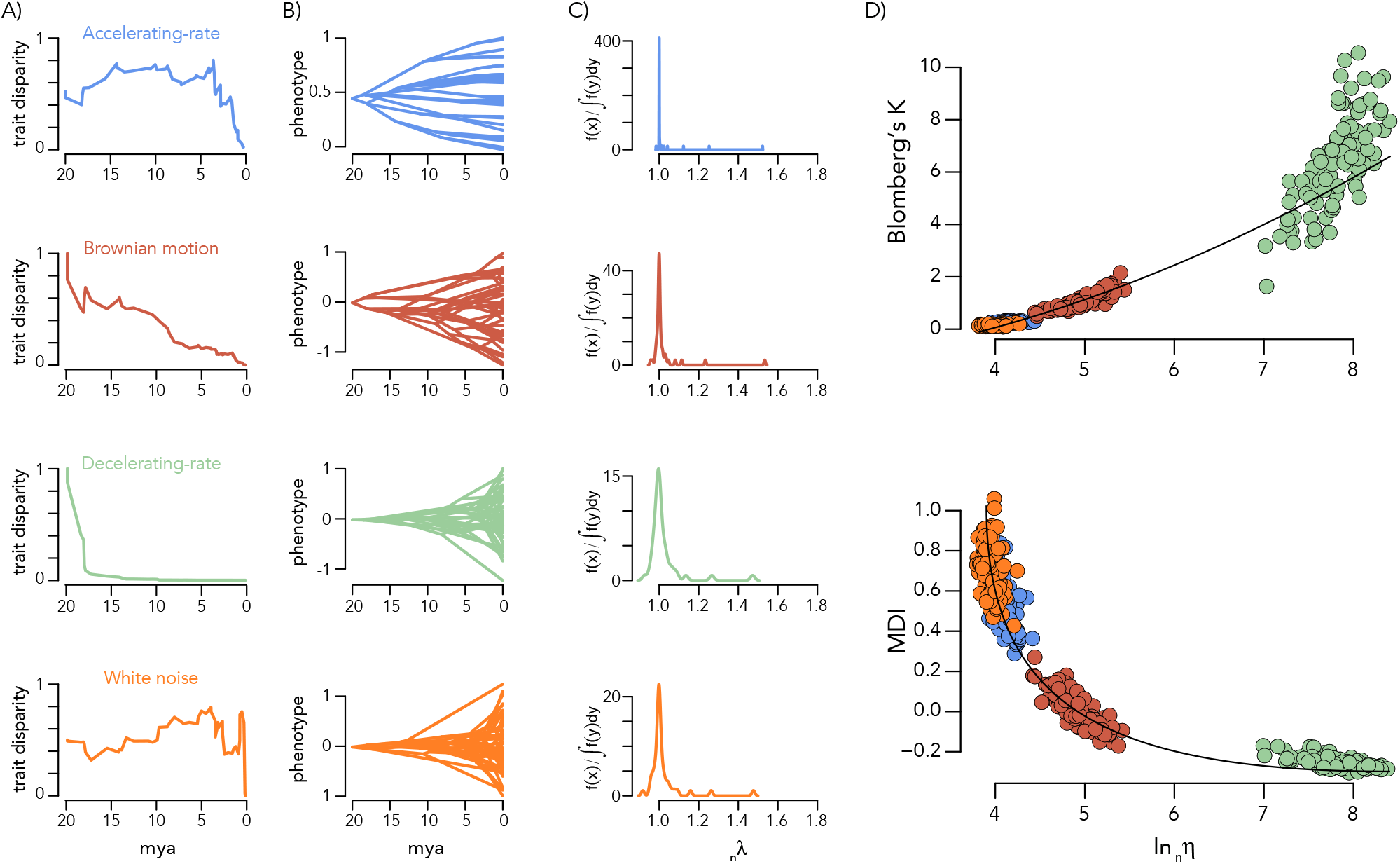
Interpreting the tracer of spectral density profiles. (A) Disparity-through-time plots for traits evolved under an AC, BM, DC, and white-noise model on the same 100-tip constant-rate birth-death phylogeny. (B) Traitgrams and (C) spectral density profiles for the phylogenetic traits in (A). Note the difference y-axis range in (C). Pairwise plots of Blomberg’s *K* and MDI as a function of the tracer for phylogenetic trait data simulated under AC, BM, DC, and white-noise models. The best-fit regression slopes are shown for each plot.

The fragmenter summary statistic represents the relative abundance of small *versus* large _*n*_λ. In macroevolutionary terms, larger *_n_ψ* indicate a more discrete distribution of trait means in trait space. We show that for trait data simulated on increasingly discretized macroevolutionary landscapes, spectral density profiles have correspondingly higher fragmenter values (Fig. 4A-C). We also show for trait data simulated with DC models with an increasingly negative rate parameter, *β*, which produce increasingly discretized trait space, that spectral density profiles have correspondingly higher fragmenter values (Fig. 4D-F). Notably, the discrete clusters of mean trait values generated by macroevolutionary landscapes are generally not monophyletic, whereas those generated by DC models are monophyletic.

**Figure 4:**
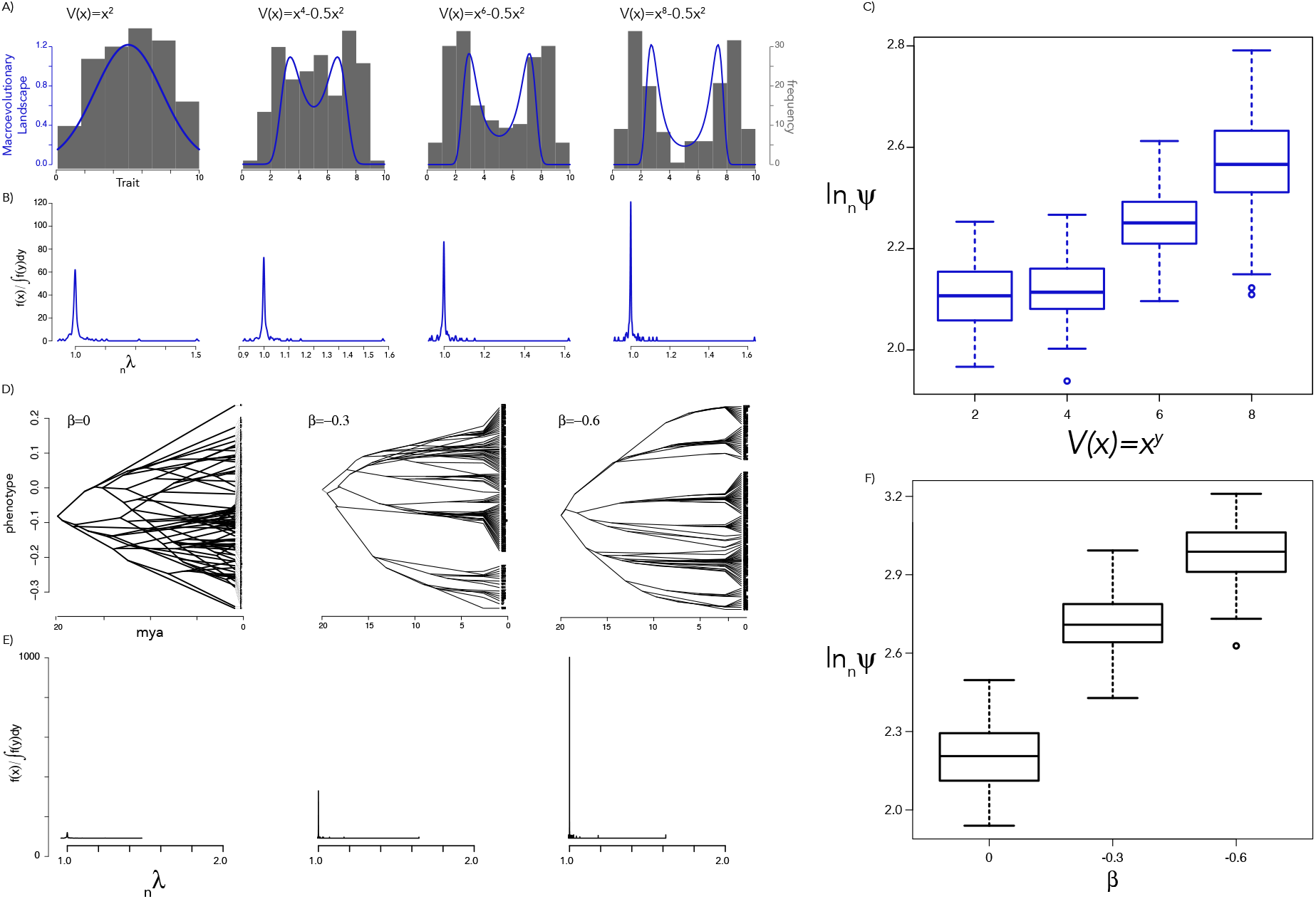
Interpreting the fragmenter of spectral density profiles. (A) Histograms of simulated trait values (grey) under four macroevolutionary landscapes (blue). (B) Spectral density profiles for phylogenetic trait data simulated under each landscape in (A). (C) boxplot of fragmenter values for spectral density profiles generated under each macroevolutionary landscape in (A). (D) Traitgrams of phylogenetic trait data simulated under ACDC models with different rate parameter values, *β*. (E) Spectral density profiles for the phylogenetic trait data in (D). (F) Boxplot of fragmenter values for phylogenetic trait data simulated under each DC model in (C).

The splitter summary statistic, which is the principal _*n*_λ computed from the nMGL, is diagnostic of the bipartiteness of the nMGL. Specifically, it is indicative of how easily the graph can be disjointed into two components. We show that splitter values increase (approaches 2) as the number of monophyletic groups with different trait means approaches two (Fig. 5A-C). When groups are defined using k-means clustering (with *k* = 2) on the nMGL, the average phylogenetic distance between groups approaches two times the crown age of the phylogeny when there are two monophyletic groups, demonstrating that clustering on the nMGL allows recovering these two groups (Fig. 5D). Splitter values obtained from the randomized datasets are similar to those obtained from the original datasets, suggesting that phylogenetic signal has little effect on splitter values (Supplemental Fig. 1).

**Figure 5:**
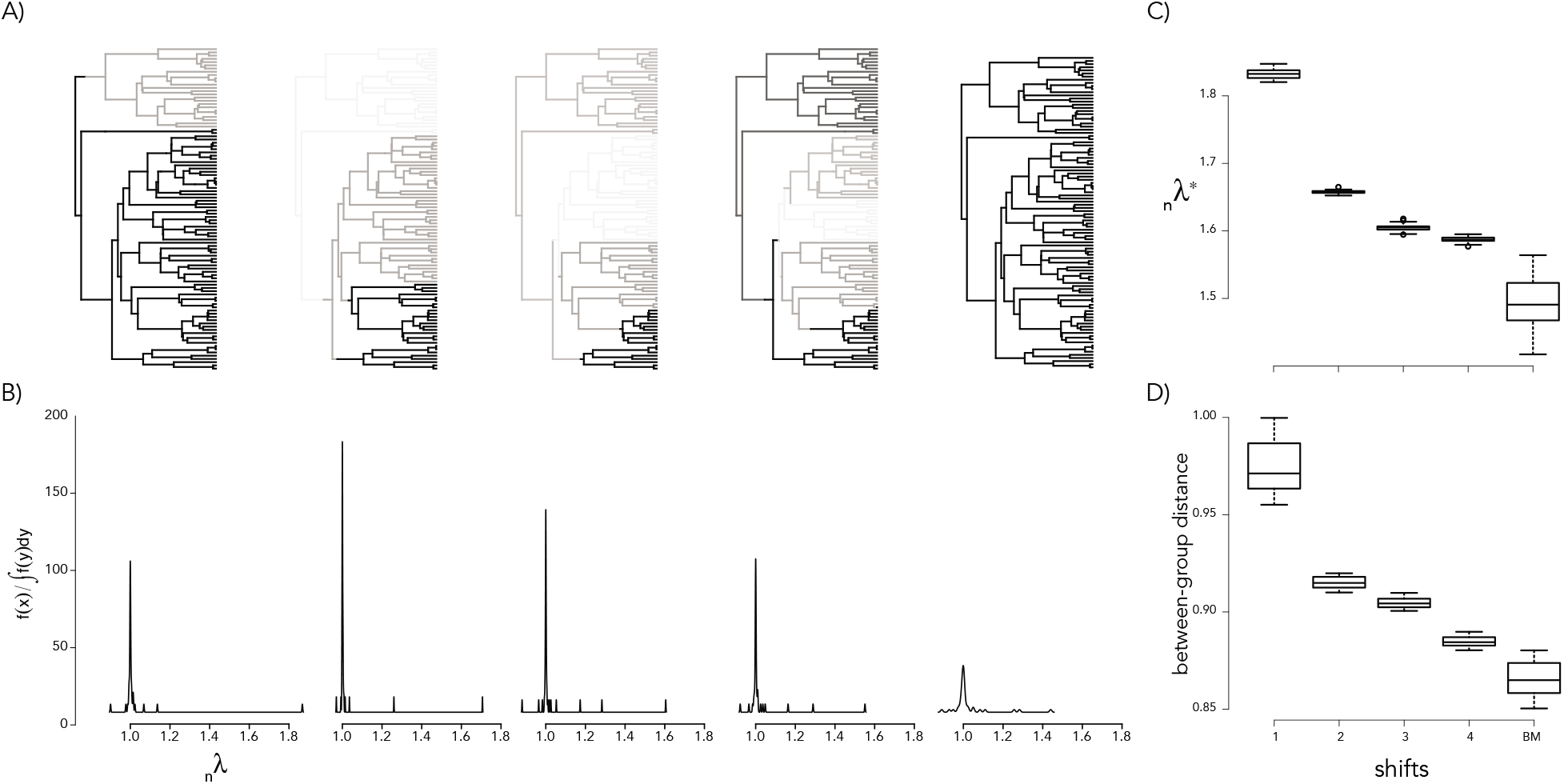
Interpreting the splitter of spectral density profiles. (A) Phylogenies simulated with 1 – 4 monophyletic shifts in mean trait values and no shifts in trait value. Different mean trait values are represented in grey scale. (B) Spectral density plots for the phylogenetic trait data in (A). (C) Boxplot of splitter values for phylogenetic trait data simulated under different numbers of monophyletic shifts in mean trait value. (D) Boxplot of the between-cluster branch-length distances (as a ratio over two times the crown age of the tree) for phylogenetic trait data simulated under different shifts in mean trait value, where clusters are defined by k-means clustering on nMGL (with k=2). Both splitter and between-cluster branch-length distance increase as the nMGL approaches bipartiteness (splitter=2).

Importantly, the fragmenter and tracer values are sensitive to the introduction of erroneous data, although not dramatically (Supplemental Fig. 2). When a considerable amount of sampling variance (equal to three times the standard error) is introduced on 10% of tips, fragmenter and tracer values decrease only slightly. The impact of erroneous data only becomes appreciable when it is introduced on a large proportion of tips (≥ 30%). However, it is only when 100% of tips are affected by erroneous data that the inference of fragmenter and tracer values begins to approach that of AC models (*β* = 1.5), which shows that the nMGL is in large robust to error-prone data.

### Clustering Models of Phylogenetic Trait Data

For the traits simulated on the constant-rate birth-death trees under the three different trait evolution models, we found that the spectral density profiles were optimally clustered into three groups (bootstrap probability > 0.95) (Fig. 6A). Separate clusters could be overwhelmingly (> 95%) assigned to AC, BM, and DC models with an average silhouette width= 0.6. The DC cluster is considerably farther from the AC and BM clusters than the AC and BM clusters are from each other, based on Euclidean distance. Trait models simulated on increasing-rate and decreasing-rate trees show slightly different abilities to cluster trait models using spectral density profiles. They also show different configurations of profiles in multidimensional space, although this is expected because the nMGL is computed using the phylogenetic distance matrix, which is sensitive to tree shape. For the increasing-rate tree, the profiles were optimally clustered into three groups (bootstrap probability > 0.95), each of which could be exclusively assigned to either AC, BM, or DC models with an average silhouette width= 0.79 (Fig. 6B). Similarly to the constant-rate tree, the DC cluster is considerably farther from the AC and BM than the AC and BM are from each other. For the decreasing-rate tree, we found two significant clusters (bootstrap probability > 0.95), one of which can be exclusively assigned to DC models and another that is a hodgepodge of AC and BM trait models with an average silhouette width= 0.55 (Fig. 6C). When plotted in multidimensional space, the AC and BM models simulated on the decreasing-rate tree occupy the same region and are therefore indistinguishable based on their spectral density profile summary statistics for this tree.

**Figure 6:**
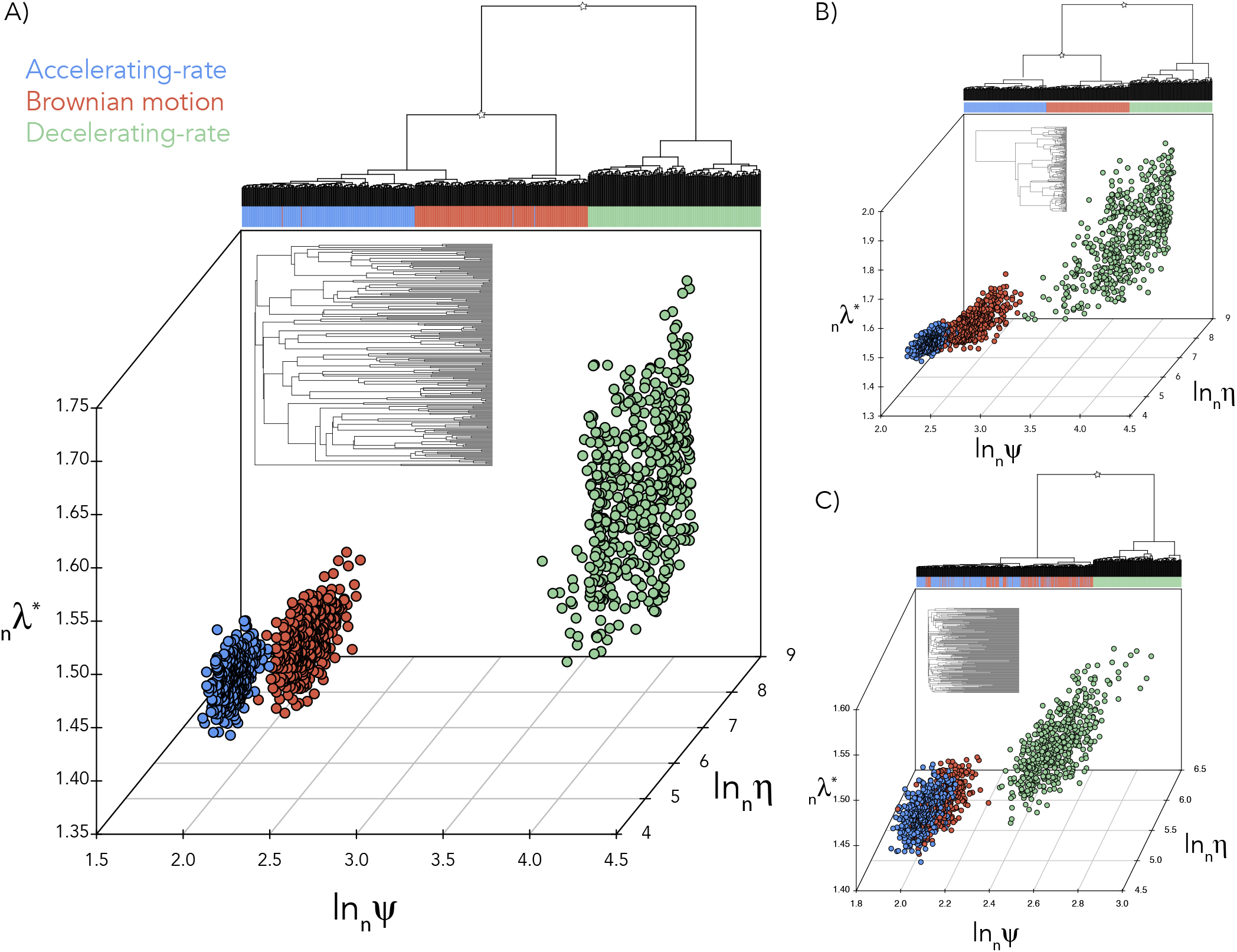
Clustering phylogenetic trait data using the spectral density profile of the nMGL. Hierarchical clustering of spectral density profiles and three-dimensional plotting of spectral density profile summary statistics for phylogenetic trait data simulated under AC, BM, and DC models of trait evolution on (A) a constant-rate birth-death tree, (B) an increasing-rate birth-death tree, and (C) a decreasing-rate birth-death tree. The trees are shown as insets. Asterisks denote bootstrap probabilities > 0.95 at the split.

For the constant-rate non-ultrametric tree, trait models are also distinguishable based on hierarchical and k-medoids clustering (Supplemental Fig. 3A). The average silhouette width for clusters of traits on the non-ultrametric tree is 0.82, compared to only 0.6 on the ultrametric tree (Supplemental Fig. 3B), which demonstrates that the trait models are more distinguishable on the non-ultrametric tree. We similarly found trait models to be distinguishable on increasing-rate (Supplemental Fig. 3C) and decreasing-rate (Supplemental Fig. 3D) non-ultrametric trees.

We estimated the effects of tree size on spectral density profile summary statistics. Fragmenter and tracer values increase with tree size, while splitter values decrease with tree size (Supplemental Fig. 4A). At 20 tips, the profiles of AC, BM, and DC models occupy the same phylogenetic trait space, but at 50 tips the models are distinguishable (Supplemental Fig. 4B). While the nMGL loses some information on the size of the graph compared to the non-normalized version, clearly there is still some effect of size. This is likely because size and shape are integrated in phylogenies.

### Applications

Traditionally, likelihood-based models are fit to phylogenetic trait data and the model showing the best support is inferred as the generative one. Oftentimes the difference in support between models is small and therefore finding traits with similar evolutionary histories or comparing those evolutionary histories can be difficult. The ability of our approach to directly compare the spectral density profiles of the nMGLs of different traits on the same tree allows us to find clusters of traits with similar evolutionary histories and then compare those histories in a multidimensional space defined by interpretable parameters without needing to qualify differences based on estimated likelihoods.

By simulating datasets under different trait models on the Cetacean phylogeny, we are able to visualize how distinguishable these models are from one another under different parameter values (Fig. 7). When all parameter values are taken together, we are not able to clearly distinguish between all models using hierarchical clustering (Fig. 7). While under certain parameter values each model occupies its own space, there is nonetheless overlap for parameter values, suggesting that, for the Cetacean phylogeny, trait evolution under different phenotypic models are quite similar. Particularly similar models are DC and DD, although these diverge in phylogenetic trait space for large parameter values; and OU and AC, but unsurprisingly, because these two models are algebraically identical on ultrametric trees (Uyeda et al. 2015).

**Figure 7:**
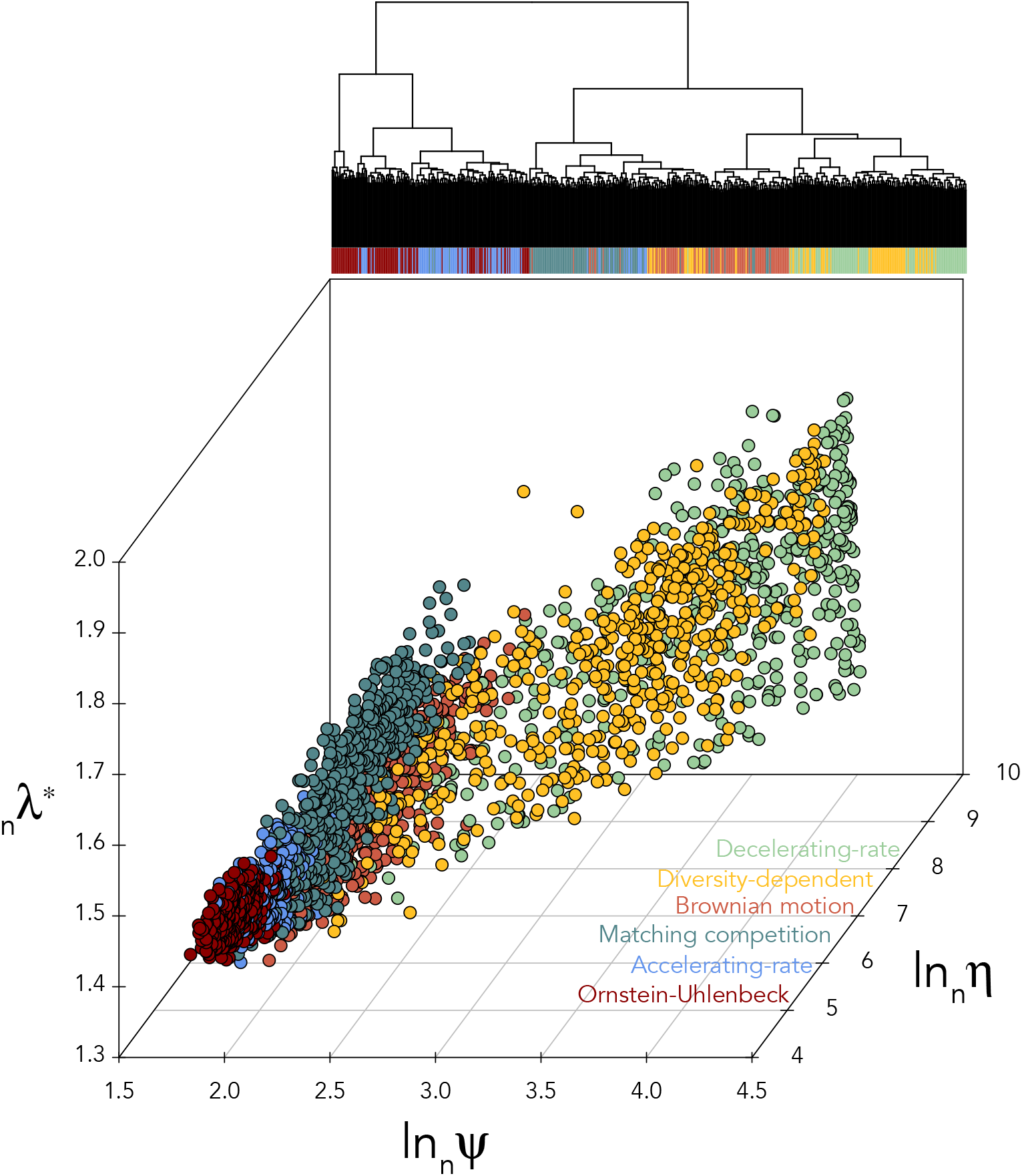
Spectral density profiles for simulated trait models on the Cetacean phylogeny. Hierarchical clustering and multidimensional plot of spectral density profile summary statistics for trait data simulated under AC, BM, DC, DD, MC, and OU models under varying parameter values on the Cetacean phylogeny.

We clustered the spectral density profiles for 27 pPC traits in the tanager phylogeny. We identified two clusters using hierarchical clustering (bootstrap probability= 0.96) (Fig. 8A); and found the same two clusters using k-medoid clustering, where the inferred axes explained 69% of variance among the spectral density profiles (Fig. 8B). Cluster 1 was comprised of 10 plumage traits, 6 resource-use traits, and 1 song trait, whereas Cluster 2 was comprised of 9 song traits and 1 resource-use trait, suggesting different evolutionary histories for different types of traits (Fig. 8B). Cluster 1 showed significantly higher (*T* > 2.8, *P* < 0.01) splitter, fragmenter, and tracer values compared to Cluster 2 (Fig. 8C). This suggests that that plumage and resource-use traits have a stronger phylogenetic signal and evolve into more discrete trait space, indicative of monophyletic clusters of traits. While the plumage and resource-use cluster have a significantly higher splitter value than the song cluster, both have low splitter values (i.e., ≪ 2) and therefore little evidence of bipartiteness.

**Figure 8:**
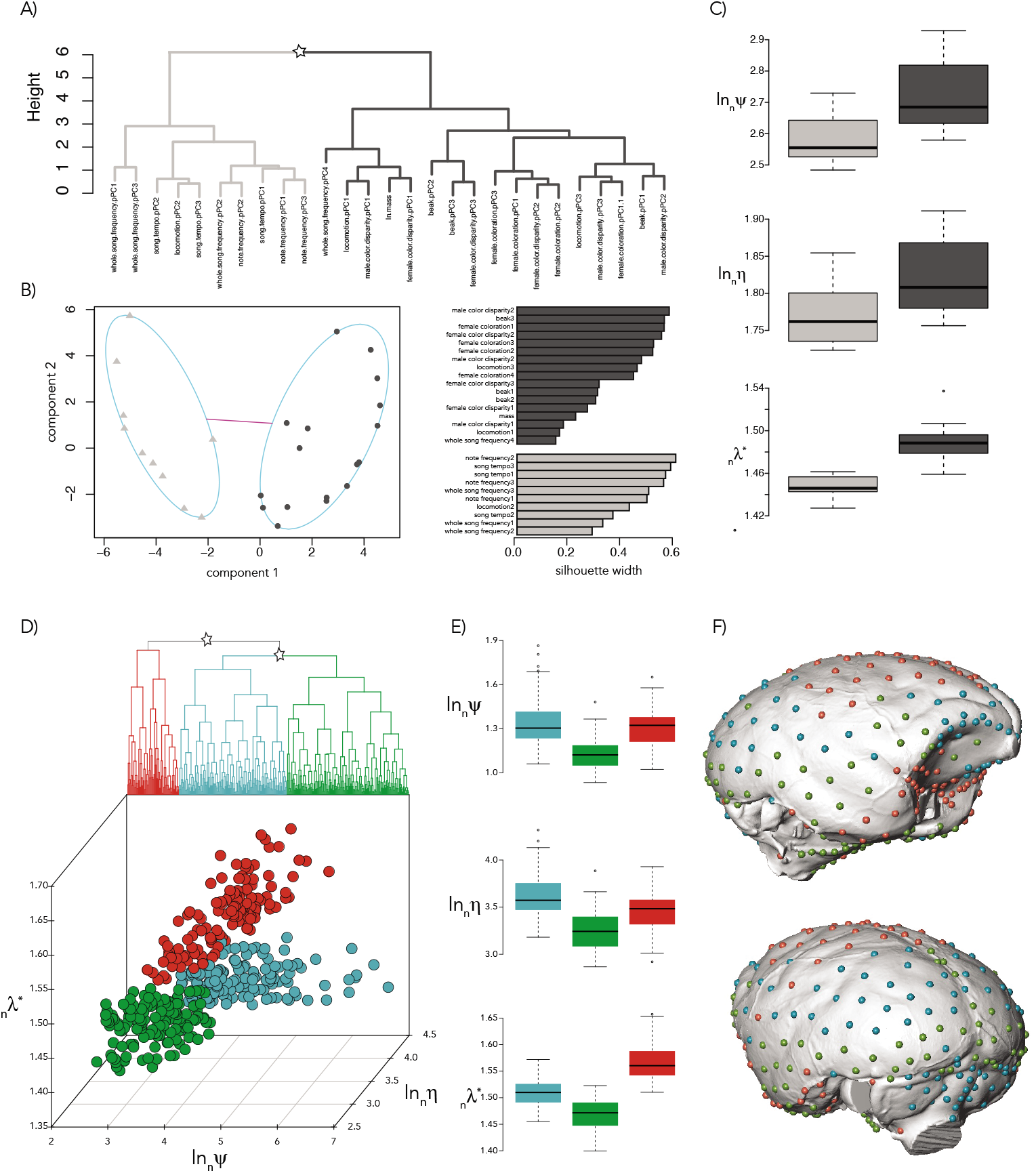
Spectral density profiling of traits in tanagers and New World monkeys. (A) Hierarchical and (B) k-medoids clustering on the spectral density profiles of the nMGLs constructed from 27 pPC traits on the tanager phylogeny. Silhouette widths are shown for each pPC trait in the k-medoid clustering. (C) Spectral density profile summary statistics for pPC traits within each cluster identified in (A,B). (D) Hierarchical clustering of spectral density profiles and multidimensional plot of spectral density profile summary statistics for 399 landmarks on New World monkey endocrania: cluster 1 (blue), cluster 2 (green), cluster 3 (red). (E) Boxplot of summary statistics for each cluster identified in (D). (F) Threedimensional representation of the New World monkey endocranium with placement of the clusters of landmarks corresponding to (D).

The landmark data for New World monkeys clustered into three groups according to k-medoid clustering, with a minimum average silhouette width of 0.51, and according to hierarchical clustering (bootstrap probability> 0.9) (Fig. 8D). Cluster 1 showed significantly higher (*T* > 1.96, *P* ≤ 0.05) fragmenter and tracer values, suggesting a stronger phylogenetic signal and evolution into more discrete trait space compared to the other clusters of landmarks (Fig. 8E). Cluster 2 showed significantly lower fragmenter and tracer values than the other clusters, suggesting it evolves with little phylogenetic signal into a more uniform trait space. Cluster 3 showed intermediary fragmenter and tracer values, but significantly higher splitter values, indicative of more bipartiteness. The relationship between fragmenter and tracer, which is indicative of the amount of convergence in trait space, shows that tracer values increase as a function of fragmenter values faster based on a one-sided t-test (*P* < 0.05) in cluster 1 compared to clusters 2 and 3, suggesting lower levels of convergence in cluster 1. Interestingly, the three clusters broadly correspond to well-defined brain regions (Fig. 8F). Specifically, cluster 1 comprises landmarks mostly located on the parietal, cerebellar, and the anterior portion of the frontal region, cluster 2 landmarks mainly correspond to the temporal, occipital, and stem regions, and cluster 3 comprises landmarks on the posterior and ventral areas of the frontal region and parts of the temporal. These results suggest that different brain regions evolved with different evolutionary histories. When we clustered the landmark data along each axis separately, treating the coordinates as tip data, we identified the same three clusters along each axis according to k-medoid clustering, with a minimum average silhouette width for each cluster of 0.46, and according to hierarchical clustering (bootstrap probability> 0.9).

## Discussion

We recently introduced an approach for characterizing and comparing phylogenies using the spectrum of the graph Laplacian (Lewitus and Morlon 2016a). Here, we have extended this approach to analyse the evolution of traits within phylogenetic clades. We have shown how to compute the spectral density profile of the nMGL for phylogenies with associated trait data and demonstrated how to use these profiles to characterize and compare trait data within a phylogenetic clade. This provides a broad, scalable framework for characterizing the distribution of traits within a phylogenetic clade without classifying those distributions according to pre-defined models of phenotypic evolution. This non-parametric approach therefore provides a complement to existing model-based approaches to studying phenotypic evolution.

Because the spectral density profile of the nMGL is computed directly from the phylogeny and trait data, it provides a comprehensive rendering of the structure of trait evolution across a phylogenetic clade. Consequently, the spectral density profiles of different traits on a phylogenetic tree, unlike likelihood values or summaries of phylogenetic signal, can be clustered absent any *a priori* model specification. We show that this is successful in distinguishing between phylogenetic trait data generated under different macroevolutionary processes and sensitive to the parameter values under which those processes are generated. Hence, in the same way that spectral density profiles have been used for identifying principal patterns of diversification in vertebrates (Lewitus and Morlon 2016b), they can be used for identifying principal patterns of phenotypic evolution across multiple traits within clades, as we have illustrated here with two empirical datasets. Spectral density profiles can also be used to quickly evaluate how distinguishable different trait evolutionary processes are, as we have illustrated here on the Cetacean phylogeny. This can be very useful when developing new models, to make sure they will be distinguishable before putting all the effort into develop likelihood-based inferences for these models. Similarly, although it is impossible to separate the relative contribution of phylogenetic and trait distances on the SDP, it is possible to compare SDPs for the same trait data across multiple versions of a phylogeny (e.g., a posterior distribution of trees generated by Bayesian inference) and thus estimate the effect of tree construction on inferences of trait evolution. We can also anticipate that spectral density profiles will be useful to compute the distance between simulated and real data in Approximate Bayesian Computation approaches (Beaumont 2010) for fitting models of phenotypic evolution that are not amenable to likelihood computation (e.g., Clarke et al. (2017)). Although currently limited to the analysis of continuous traits, an extension of the nMGL to incorporate discrete binary traits would be straightforward: the trait distance between species would be 0 or 1 if pairs have the same or a different trait, respectively. Existing work on signed graph Laplacians (Kunegis et al. 2010), which attach a positive, negative, or neutral sign to each edge, already show the potential for using graph Laplacians to explore graphs with associated discrete values. As there are a wide range of discrete traits that are the focus of many macroevolutionary questions (e.g., Beaulieu et al. (2013)). we think development of the nMGL for the analysis of discrete trait evolution is an important direction for future work to move in.

When reduced to their constituent properties (i.e., splitter, fragmenter, and tracer values), spectral density profiles are useful in summarizing the structure of phylogenetic trait data and in visualizing differences between them. The tracer is a measure of phylogenetic signal and correlates well with conventional summary statistics. Blomberg’s *K*, as a measure of the partitioning of within- *versus* among-clade variance, resembles what tracer is measuring, which is the iteration of _*n*_λ around a single value. When within-clade variance is low and among-clade variance is high, then the majority of _*n*_λ will have a similar value, the tracer will be high and so will Blomberg’s *K*. The fragmenter measures the discreteness of phenotypic space. Higher fragmenter values indicate that trait values are distributed in more discrete groups in phenotypic space, as would occur under an early burst model of trait diversification or high levels of convergence to multiple optima. The relationship between the tracer and fragmenter gives some indication as to whether convergence has likely occurred: the ratio of tracer to fragmenter will be higher if the discretization of trait values in phenotypic space shows a strong phylogenetic signal (i.e., in the absence of convergence). We show, for example, that a two-peak macroevolutionary landscape results in high fragmenter values, but relatively lower tracer values than occur under a DC model, indicative of the high level of phenotypic convergence in the macroevolutionary landscape model and low level of phenotypic convergence in the DC model. Of course, we cannot assign a threshold value for convergence, above which the tracer to fragmenter ratio conclusively evinces phenotypic convergence. However, for a given analysis of different trait data on a tree, we recommend comparing tracer to fragmenter ratios between analyses, in order to deduce the comparative levels of convergence between datasets. Finally, the splitter of the nMGL is diagnostic of the bipartiteness of the graph and therefore, in terms of phylogenetic trait data, higher splitter values indicate a bimodal distribution of trait values with high phylogenetic signal.

We analyse a previously published dataset on pPCs for tanagers (Drury et al. 2018) to show the usefulness of clustering phylogenetic trait data to identify and characterize traits with similar evolutionary histories among a set. Our result, that the evolution of song-related traits is distinct from that of plumage- and resource-use-related traits, is consistent with those found in Drury et al. (2018) for species that are year-round territorial and/or found in dense habitats. The high tracer and fragmenter values in plumage and resource-use traits suggests the discretized trait space of these traits possesses a high phylogenetic signal, while the low tracer and fragmenter values in song traits suggests low phylogenetic signal and non-discretized trait space.

We analyse a dataset of 399 landmarks on the endocrania of 48 species of New World monkeys. We show that these landmarks cluster into three groups. Landmarks within each cluster delineate meaningful regions of the external brain morphology, which suggests that each of these regions evolved differently. Cluster 1, which mostly represents the anterior frontal, parietal, and cerebellar regions, shows these regions have evolved into a discretized trait space with high phylogenetic signal, whereas cluster 2, which defines the temporal, occipital, and stem regions, shows these regions have evolved in a more uniform space with low phylogenetic signal. Cluster 3, which defines the posterior and ventral areas of the frontal region and part of the temporal region, also shows evidence of these regions evolving into a discretized trait space, but with higher levels of convergence than the regions of cluster 1. Despite the differences in approach, these results align well with a previous analysis of the same dataset conducted using PCA (Aristide et al. 2016), which suggests that, during the adaptive radiation of New World monkeys, brain shape evolved first into discrete regions of morphospace, with subsequent bursts of evolution generating convergence among clades. Moreover, according to Aristide et al. (2016), the different stages of this diversification can be associated to the evolution of particular regions of the brain. For example, coincident with our results for cluster 1, the anterior frontal region would have diversified early into discrete trait optima, while convergent changes would be mostly associated with other areas of the frontal region, in agreement with our cluster 3. Overall, our results support the idea that there has been differential selection on different brain regions in New World monkeys, due both to an early adaptive radiation and convergence on ecologically relevant traits (Rosenberger 1992; Gavrilets and Losos 2009; Aristide et al. 2015, 2016).

A major focus of work on phenotypic evolution relates to the study and identification of co-evolving traits using multivariate models (Clavel et al. 2015). Specifically, the correlated evolution of multiple traits resulting in evolutionary integration expects such sets of traits to have shared evolutionary histories (Goswami 2007). We would therefore also expect that these traits, whether they are biologically integrated or co-evolving with some shared variable, will have similar spectral density profiles; and so clustering profiles may be a way to identify different sets of integrated traits from multivariate data. This can become particularly useful when there are many traits, as is more often becoming the case with the proliferation of trait data (e.g., Jones et al. (2009); Hamish et al. (2014)).

We have developed an approach, implemented in user-friendly software, which is a valuable addition to existing PCMs and provides a new way to analyse and conceive phenotypic evolution.

## Acknowledgments

We would like to thank Alex Bjarnason, Julien Clavel, Jonathan Drury, Carmelo Fruciano, Odile Maliet, Marc Manceau, Olivier Missa, and Guilhem Sommeria Klein for helpful comments on the manuscript. EL would like to thank Evan Charles for helpful discussion. Funding was provided by the CNRS and a grant (PANDA) from the European Research Council (ERC) attributed to HM. The views expressed are those of the authors and should not be construed to represent the positions of the U.S. Army, the Department of Defense, or the Department of Health and Human Services.

---

**Figure S1:**
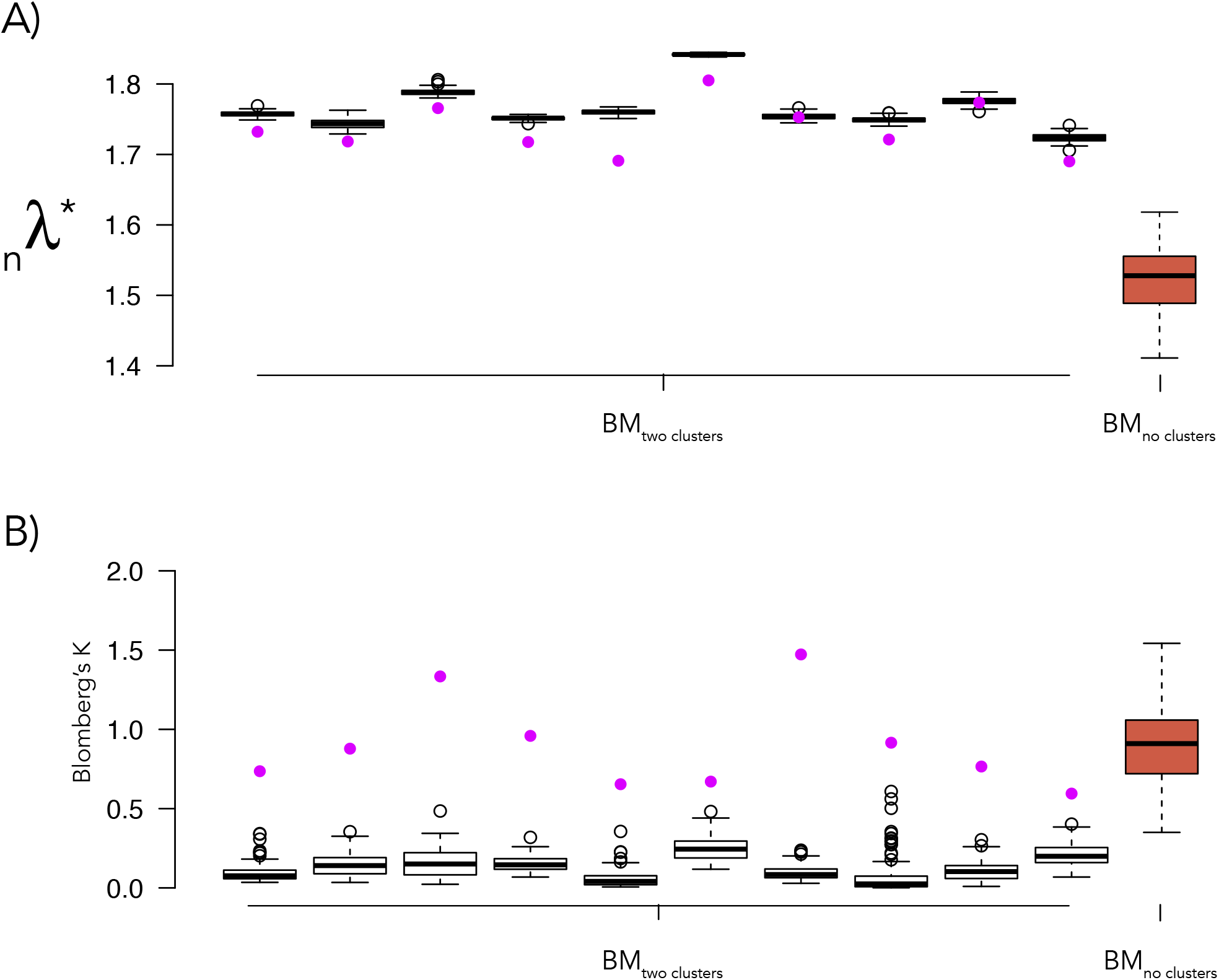
Measuring the effect of phylogenetic signal on splitter values. (A) Boxplot of the splitter values for 100 randomized datasets (white) obtained for each of the ten datasets with two monophyletic clusters. Splitter values for the initial BM datasets with two clusters are shown in purple. Boxplot of 100 datasets simulated under a simple BM process with no clusters on a single tree (coral) is shown for comparison. (B) Boxplot of Blombergs *K* for each randomized dataset (white); values for the initial BM datasets with two clusters are shown in purple. Boxplot of 100 datasets simulated under a simple BM process with no clusters on a single tree (coral) is shown for comparison.

**Figure S2:**
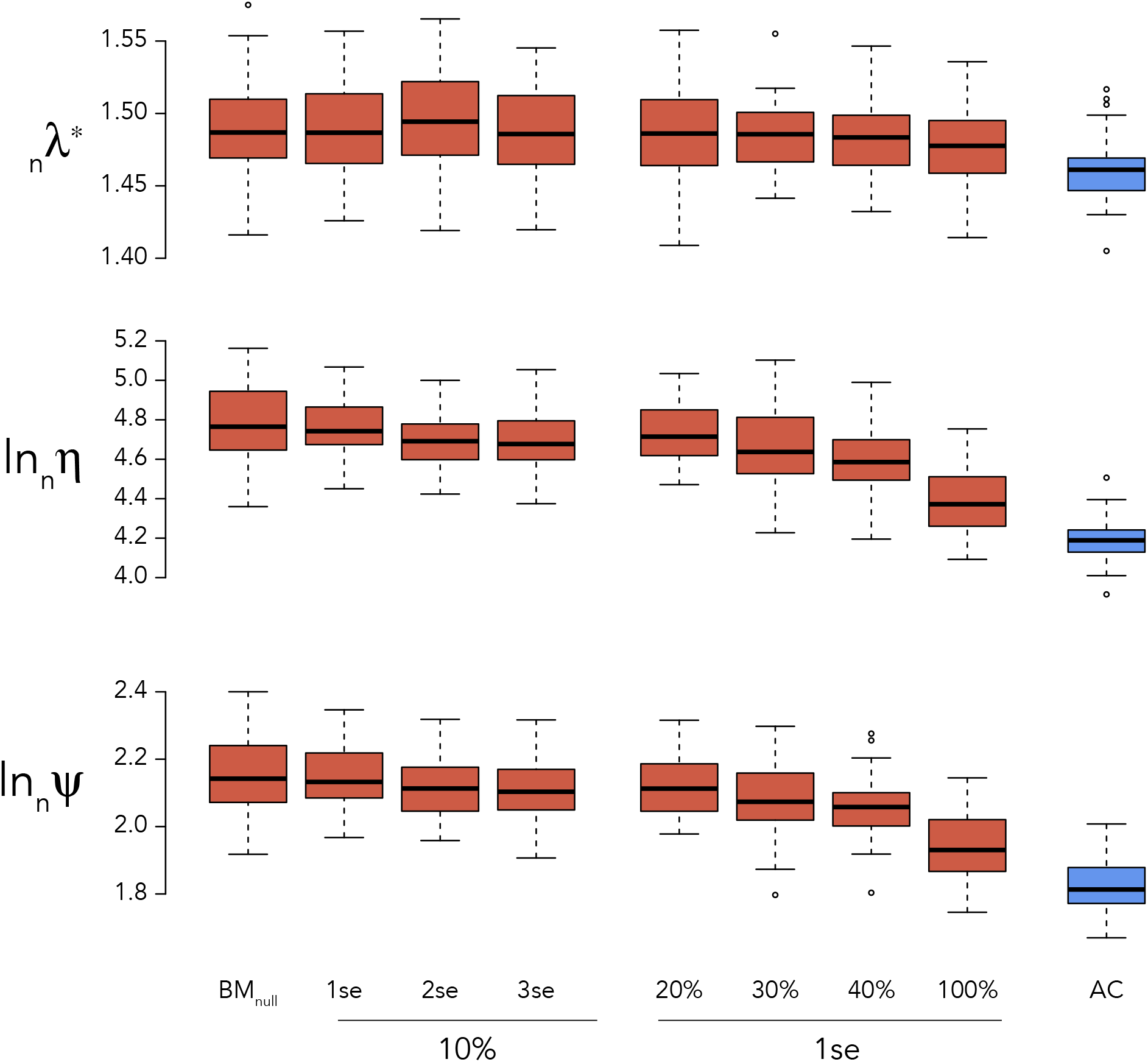
Measuring the effect of erroneous trait data on spectral density profile summary statistics. Spectral density profile summary statistics for data simulated under a BM process (coral) with introduced error for 10% of tips with a sampling variance equal to one, two, and three times the standard error of the simulated BM data; and with a sampling variance equal to one times the standard error for 10, 20, 30, 100% of tips. Spectral density profile summary statistics for data simulated on the same tree under an ACDC process (*β* = 1.5) is also shown (cornflowerblue).

**Figure S3:**
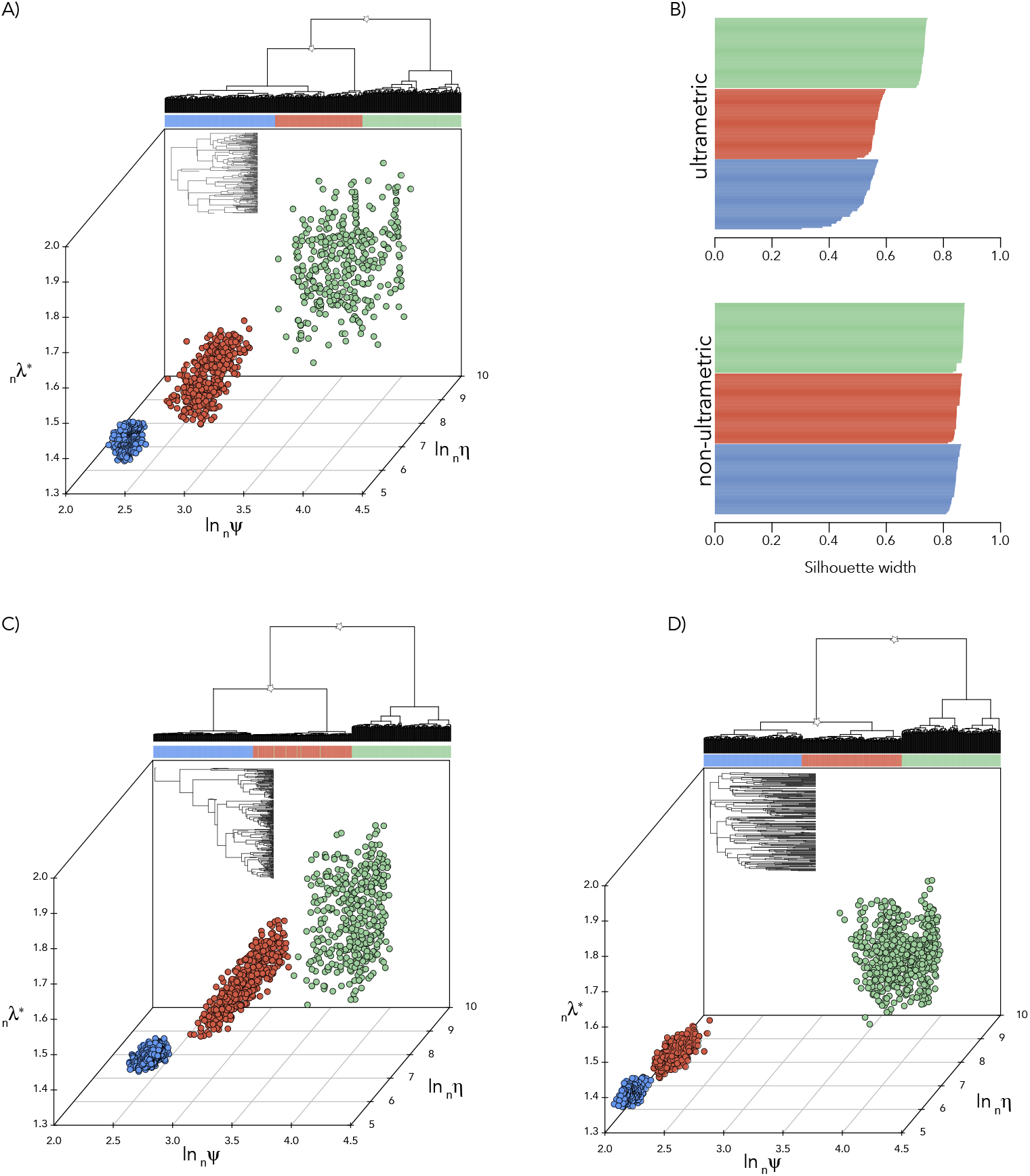
Clustering phylogenetic trait data using the spectral density profile of the nMGL on a non-ultrametric tree. Hierarchical clustering of spectral density profiles and three-dimensional plotting of spectral density profile summary statistics for phylogenetic trait data simulated under AC (cornflower blue), BM (coral), and DC (sea green) models of trait evolution on a single (A) constant-rate, (C) increasing-rate, and (D) decreasing-rate birth-death tree without pruning extinct lineages. Tree is shown in inset. Asterisks denote bootstrap probabilities > 0.95 at the split. (B) Silhouette widths for profiles comprising each trait model cluster simulated on the ultrametric or non-ultrametric tree (see Fig. 5A).

**Figure S4:**
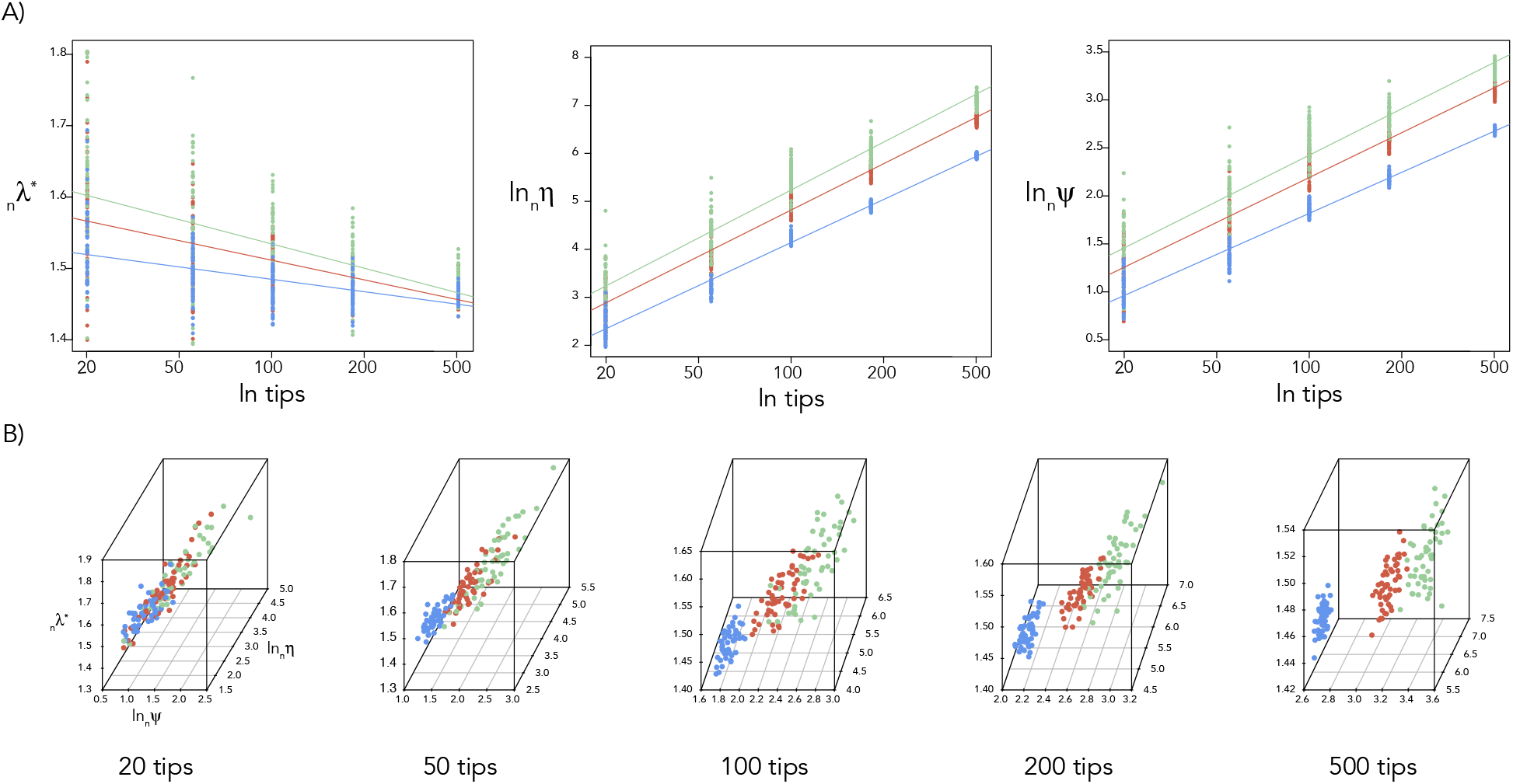
Effect of tree size on the nMGL. (A) Scatterplots and OLS regression slopes for spectral density profile summary statistics for trait data simulated under DC (sea green), BM (coral), and AC (cornflower blue) models on constant-rate birth-death trees with different numbers of tips. (B) Phylogenetic trait space for trait models simulated under AC, BM, and DC models on trees with different numbers of tips.

